# Spike substitution T813S increases Sarbecovirus fusogenicity by enhancing the usage of TMPRSS2

**DOI:** 10.1101/2023.01.15.524170

**Authors:** Yong Ma, Pengbin Li, Yunqi Hu, Tianyi Qiu, Lixiang Wang, Hongjie Lu, Kexin Lv, Mengxin Xu, Jiaxin Zhuang, Xue Liu, Suhua He, Bing He, Shuning Liu, Lin Liu, Yuanyuan Wang, Xinyu Yue, Yanmei Zhai, Wanyu Luo, Haoting Mai, Wenjing Zhao, Jun Chen, Shoudeng Chen, Xiaoli Xiong, Mang Shi, Ji-An Pan, Yao-Qing Chen

**Affiliations:** School of Public Health (Shenzhen), Sun Yat-sen University, Shenzhen, Guangdong Province, China; Institute of Clinical Science, Zhongshan Hospital, Shanghai Medical College, Fudan University, Shanghai, China; The Center for Infection and Immunity Study and Molecular Cancer Research Center, School of Medicine, Shenzhen Campus of Sun Yat-sen University, Shenzhen, Guangdong Province, China; State Key Laboratory of Respiratory Disease, CAS Key Laboratory of Regenerative Biology, Guangdong Provincial Key Laboratory of Stem Cell and Regenerative Medicine, Guangzhou Institutes of Biomedicine and Health, Chinese Academy of Sciences, Guangzhou, Guangdong Province, China; Department of Immunology and Microbiology, Zhongshan School of Medicine, Sun Yat-sen University, Guangzhou, Guangdong Province, China; National Medical Products Administration Key Laboratory for Quality Monitoring and Evaluation of Vaccines and Biological Products, Sun Yat-sen University, Guanzhou, Guangdong Province, China; Molecular Imaging Center, Guangdong Provincial Key Laboratory of Biomedical Imaging, the Fifth Affiliated Hospital, Sun Yat-sen University, Zhuhai, Guangdong Province, China

## Abstract

SARS-CoV Spike (S) protein shares considerable homology with SARS-CoV-2 S, especially in the conserved S2 subunit (S2). S protein mediates coronavirus receptor binding and membrane fusion, and the latter activity can greatly influence coronavirus infection. We observed that SARS-CoV S is less effective in inducing membrane fusion compared with SARS-CoV-2 S. We identify that S813T mutation is sufficient in S2 interfering with the cleavage of SARS-CoV-2 S by TMPRSS2, reducing spike fusogenicity and pseudoparticle entry. Conversely, the mutation of T813S in SARS-CoV S increased fusion ability and viral replication. Our data suggested that residue 813 in the S was critical for the proteolytic activation, and the change from threonine to Serine at 813 position might be an evolutionary feature adopted by SARS-2-related viruses. This finding deepened the understanding of Spike fusogenicity and could provide a new perspective for exploring *Sarbecovirus*’ evolution.

**Author Summary:** The Spike strain of SARS-CoV-2 has accumulated many mutations during its time in circulation, most of which have occurred in the S1 region, and more specifically in the RBD, in an effort to either improve the virus’s affinity for the receptor ACE2 or to enhance its ability to evade the immune system. Mutations in the Spike S2 region have more far-reaching effects than those in the S1 region because it is more conserved across sarbecoviruses. By comparing SARS and SARS2, we found that an important substitution at amino acid position 813 in the S2 region (T813S) disrupts the utilization of TMPRSS2 and can significantly influence viral entry into cells. This discovery deepens our knowledge of S proteins and provides new prospects for tracing the evolution of Sarbecoviruses.

## Introduction

Since its outbreak, SARS-CoV-2 (SARS2) has caused hundreds of millions of illnesses and fatalities globally, making it by far the worst public health catastrophe of the 21st century [1]. SARS2 was the seventh human coronavirus and likely originated in a bat host [2,3], belonging to the Sarbecovirus subgenus of betacoronavirus [4,5], along with SARS-CoV (SARS), which caused an outbreak in 2002-2003 [6]. Spike (S), as the surface glycoprotein, facilitates virus’ cell entry through host receptor binding with its S1 subunit (S1) and membrane fusion mediated by its S2 subunit (S2) [7,8]. The receptor-binding domain (RBD) is located towards the C-terminus of S1 and is responsible for binding to the receptor, angiotensin-converting enzyme 2 (ACE2) [9,10]. RBD is the primary focus of vaccine designing [11–13] and neutralizing antibody screening [14–16] despite being highly genetically variable among variants [17–19]. After spike binding to ACE2, conformational change of S2 is triggered and this conformational change facilitates fusion between the viral and host cell membranes [20], this process is believed to be highly modulated by proteolysis.

Spike protein contains multiple proteolytic sites, while a single arginine is usually located at the S1/S2 boundary and susceptible to trypsin-like protease cleavage. It is unique among Sarbecovirus spikes that SARS2 S contains a multi-basic cleavage site (681-PRRA-684), between the S1 and S2 subunits [21]. It has been proposed that cleavage at the S1/S2 site facilitates S priming and promotes membrane fusion but is not essential to membrane fusion [22]. Spike S2 contains a S2’ site for cellular protease cleavage [23,24]. It has been proposed that the S2’ site must be cleaved to fully initiate the fusion process, by either transmembrane serine protease 2 (TMPRSS2) [25] on the cell surface or by cathepsin L (CTSL) [26] in the endosomes. S2 consisted of 3 key motifs for membrane fusion: fusion peptides (FPs) and heptad repeat 1 (HR1) and 2 (HR2). HR1 and HR2 form a six-helix bundle fusion core in post-fusion S [27]. FPs are pivotal in viral entry and highly conserved between β-coronaviruses [23]. However, the exact sequence of “fusion peptide” has not yet been determined [28–31]. In a previous study, two FP candidates have been identified based on cleavage sites [28], FP1 (Animo acid (AA) 788-806) located behind S1/S2 and the extremely conserved FP2 (AA815-833) located behind S2’ site.

The membrane fusion process is crucial to viral infection. Despite sharing about 76.47% amino acid identity on S protein [32], SARS and SARS2 have generated extremely distinct infection events: The SARS-CoV-2 pandemic has lasted for more than three years and the virus is likely to remain circulating. In contrast, the 2003 SARS-CoV world-wide outbreak was quickly eradicated. The exact reason for the transmissibility difference between the two related viruses is likely to be complex and yet to be understood. The ability to use ACE2 efficiently has been proposed to be a key prerequisite for viral infection [33]. It has been reported that SARS2 S binds to ACE2 with 10- to 20-fold higher affinity than SARS S [34], but another research showed similar affinities between SARS and SARS2 RBD binding with ACE2 [35]. Although receptor binding is an essential step for virus cell entry, the subsequent S2 mediated membrane fusion is also essential [36]. In this study, we investigated the fusogenic activities of S2 using a split-GFP system, and we found that SARS2 S2 mediates more robust membrane fusion than SARS S2. We further demonstrated that the threonine to serine substitution at residue 813 in S2 significantly enhance membrane fusion and probably enhance the spread of *Sarbecovirus*.

## Results

### SARS2 S2 induced more syncytia formation than SARS S2

To investigate the role of S2 in SARS2 infection, we constructed a chimera SARS2-S protein bearing SARS S2 (cSARS2-S2sars, spike2), and a chimera SARS-S protein bearing SARS2 S2 (cSARS-S2sars2, spike4) (Fig 1A). Fluorescence-activated cell sorting (FACS) experiments showed that the surface expression levels were the same between the chimeric S and their parents (Figs 1B and 1C). Western blot (WB) analysis showed no significant difference in cleavage efficiency between chimeric S and their parents with or without trypsin (Fig 1D). However, we found that the cleavage efficiency of SARS2 S was higher than that of SARS S, and trypsin treatment increased the cleavage of SARS S significantly (Fig 1D). In the following membrane fusion assays, we treated all S proteins expressed on cells with trypsin.

**Fig 1.**
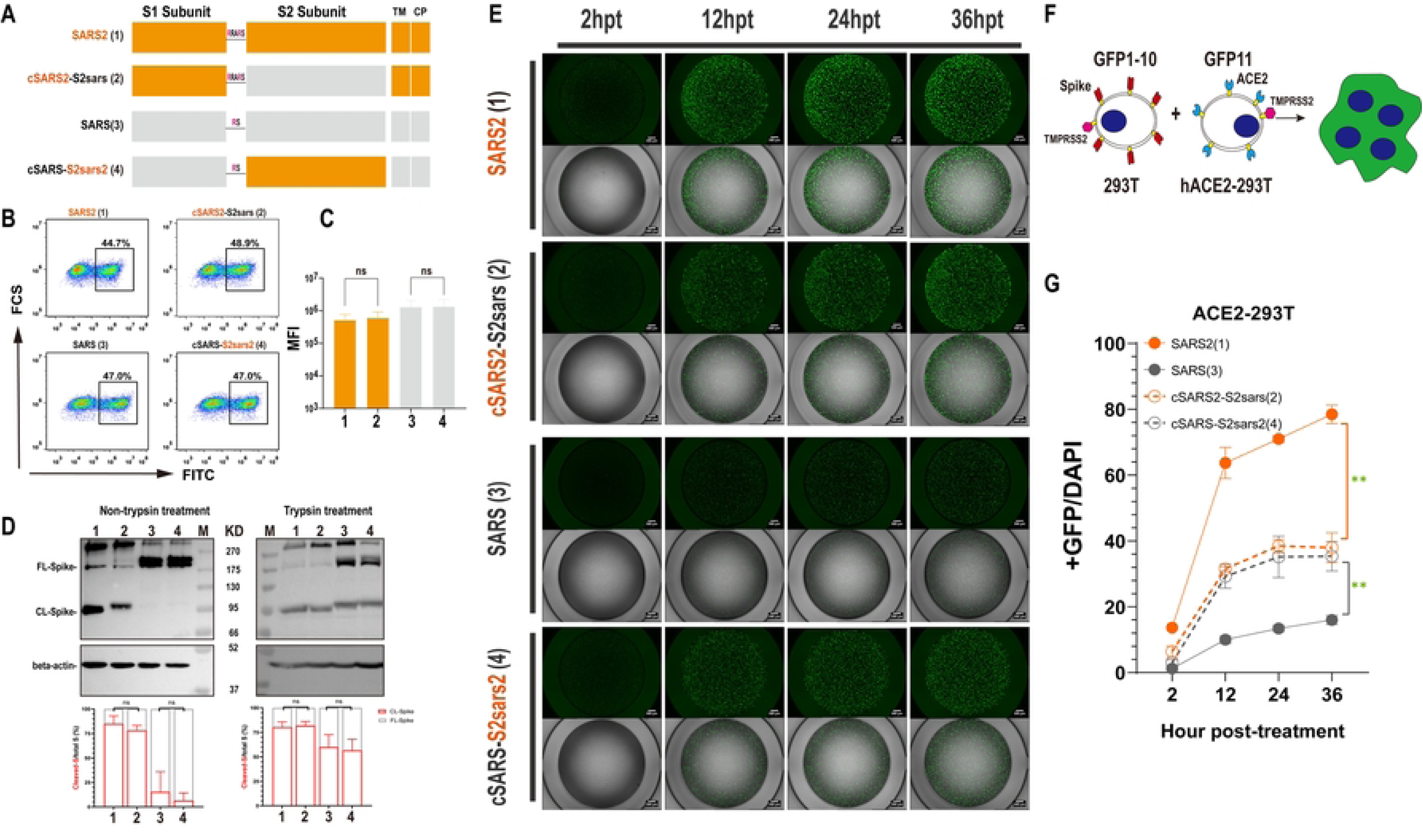
Replacement of S2 subunit affected the fusogenicity of S protein. (A) Schematic diagram for the construction of the S2-chimeric Spike. The Saffron graph depicts SARS2 S, which has the multibasic motif (RRAR) at the S1/S2 Cleavage Site; the grey graph represents SARS S. The numbers in parentheses are identical to those in Figures 1B-1G. TM, transmembrane domain; CP, cytoplasmic domain. (B and C) FACS. After transfection, the expression of surface S proteins was detected using s309 antibody which binds RBD efficiently and mouse anti-human IgG-FITC, respectively (B) and the summarized results are shown (C). MFI, mean fluorescent intensity. (D) Western blot. Representative blots of cell lysates showing spike cleavage in parental and chimeric Spikes with or without trypsin treatment. Immunoblots were probed with anti-Flag tag Abs, the full-length (FL) and cleaved (CL) S protein were marked as indicated; Beta-actin was probed as a loading control. The band intensity was densitometrically calculated using Image J, and the ratio of Cleaved-S/total S (%) was shown. (E-G) Spike-based cell-cell fusion assay. A schematic diagram showing the GFP-split system for Spike-ACE2 mediated cell fusion (F), and representative images at 2, 12, 24, 48 h post-transfection (E). The summarized results of the ratio of fusion were shown (G). Scale bar, 500 μm. Results are means +/- SD from at least three fields per condition. Results are representative of at least three independent experiments. Statistically significant differences between parental S (Spike1 or Spike3) and chimeric S (Spike2 or Spike4) were determined by a two-sided Student’s t test (C and D, ns: non-significant), or two-sided paired t test (G, *: p<0.05, **: p<0.01).

To assay the membrane fusion activities of these chimeric S proteins, we utilized a green fluorescent protein (GFP)–Split complementation system [37,38], in which GFP is split into two non-fluorescent parts, GFP1-10 and GFP11, and the reassembly of GFP1-10 and GFP11 reconstitutes a functional chromophore. With transfection, we introduced GFP1-10 and GFP11 into the donor cells transiently expressing various S proteins and the acceptor cells stably expressing ACE2 (hACE2) and TMPRSS2, respectively. The spike-mediated fusion of donor and acceptor cells could lead to the formation of syncytia. Thus, GFP1-10 and GFP11 were present in the same intracellular environment and formed a functional chromophore. The fluorescence from the chromophore reflects the fusion activities of these S proteins (Fig 1F). Two hours after the coculture of donor and acceptor cells, the cell fusion accompanied by the GFP was observed under microscopy, and the green fluorescence positive area increased rapidly with more cell fusion (Fig 1E). In this work, we quantified cell fusion activity by calculating the area ratio of GFP to DAPI. We observed that the rate and size of syncytium formation were affected by various S proteins. SARS2-S was more effective at cell fusion than SARS-S, which is consistent with their spread ability. Of note, we found the cell fusion ability of S proteins were mainly correlated with the S2. When replaced with SARS2 S2, the fusogenicity of chimeric protein cSARS-S2sars2 was remarkably improved compared with its parent, SARS; the fusion rate and size all increased; on the other side, after replacing SARS S2, the chimera S protein cSARS2-S2sars lost most of its cell fusion ability, the fusion rate and size reduced significantly (Figs 1E and 1G).

Collectively, these results suggest that S2 plays an important role in SARS2 infection, and its alternation it could affect the fusogenicity of S protein significantly.

### The Spike fusogenicity is dependent on the Internal Fusion Peptide (IFP)

To further identify key functional domains or motifs in S2 for increased spike fusogenicity, we divided S2 into three parts based on their functions [23,39] and constructed chimeric Spike5 (SARS F1, AA668-816), Spike6 (SARS F2, AA817-966), Spike7 (SARS F3, AA967-1214) and Spike10 (SARS2 F1, AA686-833), Spike11 (SARS2 F2, AA834-984), Spike12 (SARS2 F3, AA985-1213) by directly replacing amino acids of the corresponding regions on Spike1 and Spike3, respectively, as shown in Figs 2A and 2G. Via FACS and WB assay, we confirmed no significant difference in cell surface expression of these S proteins (Figs 2B and 2H); and the unchanged CL-S ratio for most chimera S, except for Spike 6, with increased cleavage; and Spike 5, with decreased cleavage (Figs 2C and 2D). Subsequent membrane fusion assays showed that swapping of F1 regions significantly affected the fusogenicity of the chimeric S proteins. The area of GFP decreased after introducing SARS F1 into SARS2-S (Spike 5, Figs 2E and 2F), while GFP signal increased significantly when SARS2 F1 was introduced into SARS1-S (Spike 10, Figs 2K and 2L). The swapping of F2 and F3 between SARS-S and SARS2-S failed to produce a similar significant alternation in the fusogenicity of S proteins as F1.

**Fig 2.**
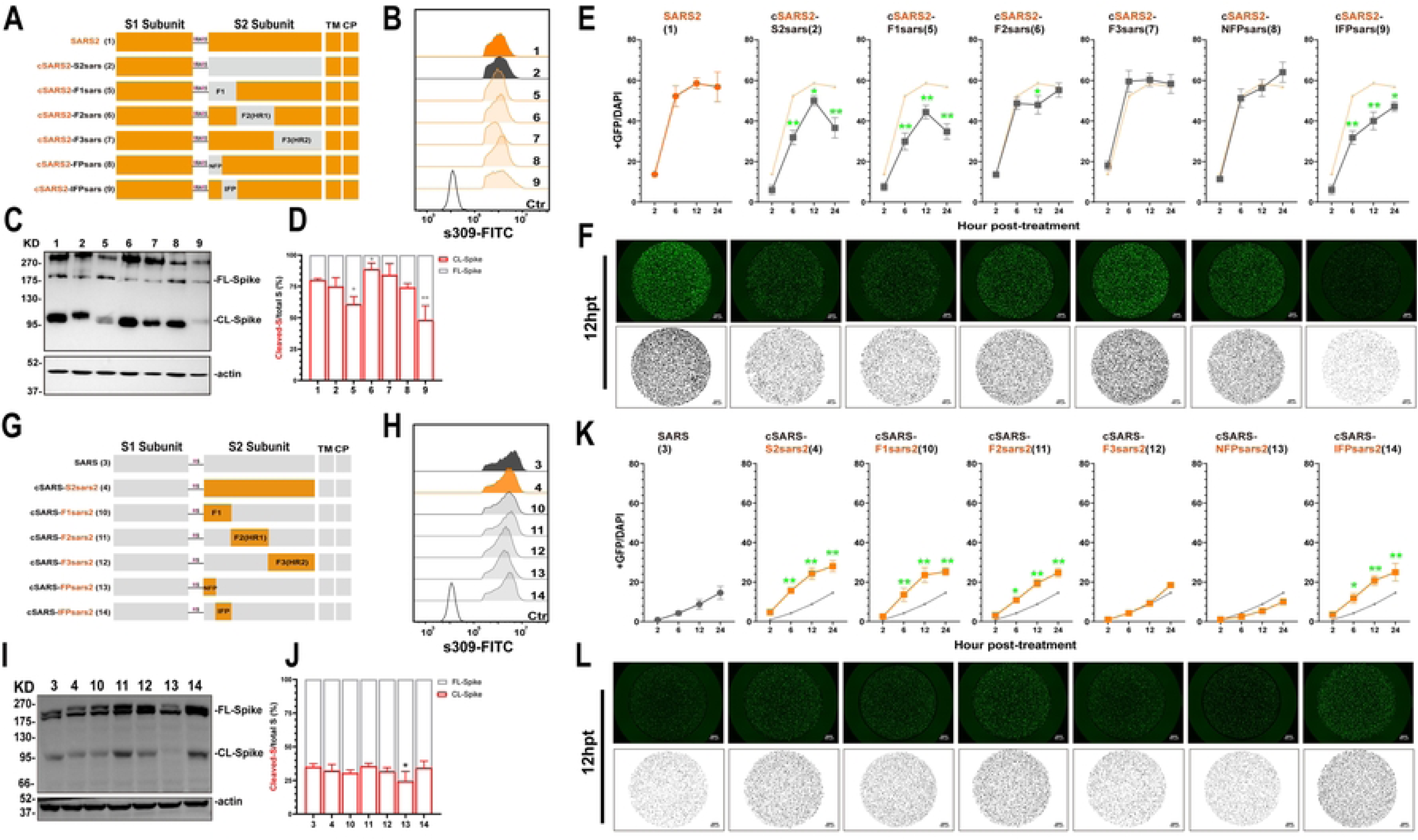
Replacing IFP motif of parental Spike influenced the fusogenicity significantly. (A and G) Schematic diagram of the S2-chimeric Spike bearing swapped motif. The S2 was divided into 3 parts according to the structure and function: F1 (686-833), F2 (834-984) and F3 (985-1170) of SARS-CoV-2 (A); F1 (655-802), F2 (803-953), F3 (954-1182) of SARS-CoV (F). Then replacing the corresponding parts with the others separately. Further, the F1 was divided into two parts: FP (686-787) and IFP (788-833) of SARS2 (A), or FP (655-756) and IFP (757-802) of SARS (F). The numbers in parentheses are identical to those in Figure 2B-2F, 2G-2L. (B and H) FACS. The summarized results of the surface S expression were shown. s309 antibody and mouse anti-human IgG-FITC were used respectively. (C, D and I, J) Western blot. A representative blot of cell lysates showing spike cleavage. FL-Spike and CL-Spike were marked as indicated. Beta-actin was used as a control. The band intensity was densitometrically calculated using Image J, and the ratio of Cleaved-S/total S (%) was shown. (E and K) Spike-based fusion assay. The fusion activity was quantified by measuring the ratio of GFP+ area to DAPI area by imaging at different times (2, 6, 12 and 24hpt). The results for SARS2, Spike4, 10-14 were shown as Saffron lines and SARS, Spike2, 5-9 were shown as grey lines, respectively. (F and L) Representative images of cell-cell fusion. Scale bar: 500 µm. Results are means +/- SD from at least three fields per condition. Results are representative of at least three independent experiments. Statistically significant differences between parental S (Spike1 or Spike3) and chimeric Spikes were determined by two-sided paired t test (D and J, *: p<0.05, **: p<0.01), or Student’s test at each point (E and K, *: p<0.05: **: p<0.01). See also Figure S1.

We further divided F1 into two portions based on whether they contained FPs or not. We define the Non-Fusion Peptide (NFP) fragment at the N-terminus and the Internal Fusion Peptide (IFP) fragment at the C-terminus. IFP fragment contained FP1 and FP2. Following the similar abovementioned strategy for swapping mutant S proteins, we constructed following chimeric S constructs. We found that after swapping NFP (Spike8, SARS AA668-769; Spike13, SARS2 AA686-787) and IFP (Spike9, SARS AA770-816; Spike14, SARS2 AA788-833), the surface expression of chimeric S was the same as the parents (Spike1 and Spike3) (Figs 2B and 2H), while the cleavage ability had a difference. We found that introducing SARS IFP into SARS2-S and introducing SARS2 NFP into SARS-S could affect the expression of S protein and decreased spike cleavage respectively (Figs 2C and 2D; Figs 2I and 2J), these results suggested direct motif replacement might affect the cleavage of S, and SARS2 FP, especially IFP, might enhance the ratio of CL-S. Subsequent membrane fusion assays showed that IFP was the key factor of S fusogenicity, replacing SARS IFP alone to SARS2 backbone (Spike9) was able to reduce S protein’s membrane fusion capacity significantly and vice versa (Spike14) (Figs 2E and 2F, Figs 2K and 2L). We also found that SARS2 F2 fragment (Spike 11) promoted fusion and this might be related to the increased S expression.

To further validate the role of IFP in S fusogenictiy, we directly replaced fragments on Spike2 and 4 for chimeric plasmid construction (S1 Figs A and F) and further assayed cell membrane fusion ability. We found that the surface expression (S1 Figs C and H) and the CL-S ratio (S1 Figs B and G) of most chimera S was the same, except for the reduction in Spike20 and the enhancement of Spike19 and 22 (S1 Figs B and G), this results further proved SARS2 IFP enhanced the cleavage of S protein. The membrane fusion results showed that introducing SARS2 F1 or IFP moderately enhanced the fusion ability of Spike2 (S1 Figs D and E), but substitution of SARS F1 and IFP significantly inhibited the fusion ability of Spike4 (S1 Figs I and J), these results confirmed that the backbone of chimeric protein might affect S fusogenicity, but IFP was the importance.

Collectively, these results showed that although there was some variation across chimera S, IFP was the most critical factor affecting fusion.

### IFP S813T mutation reduced the cell membrane fusion ability of SARS-CoV-2 S protein significantly

The IFP sequence is relatively conserved between SARS S and SARS2 S, with only six different amino acid. To investigate whether these amino acids influence the fusogenicity of S protein, we investigated the impact of each amino acid on cell membrane fusion by introducing mutations on each of them in Spike9 (Figs 3A and 3F). WB and FACS assays revealed no significant differences in expression and cleavage of each mutant S protein, except for Spike29, which is poorly expressed (Figs 3B and 3C). Subsequent membrane fusion assays revealed that only the T813S mutation significantly increased the chimeric protein’s fusion capability (Figs 3D and 3E).

**Fig 3.**
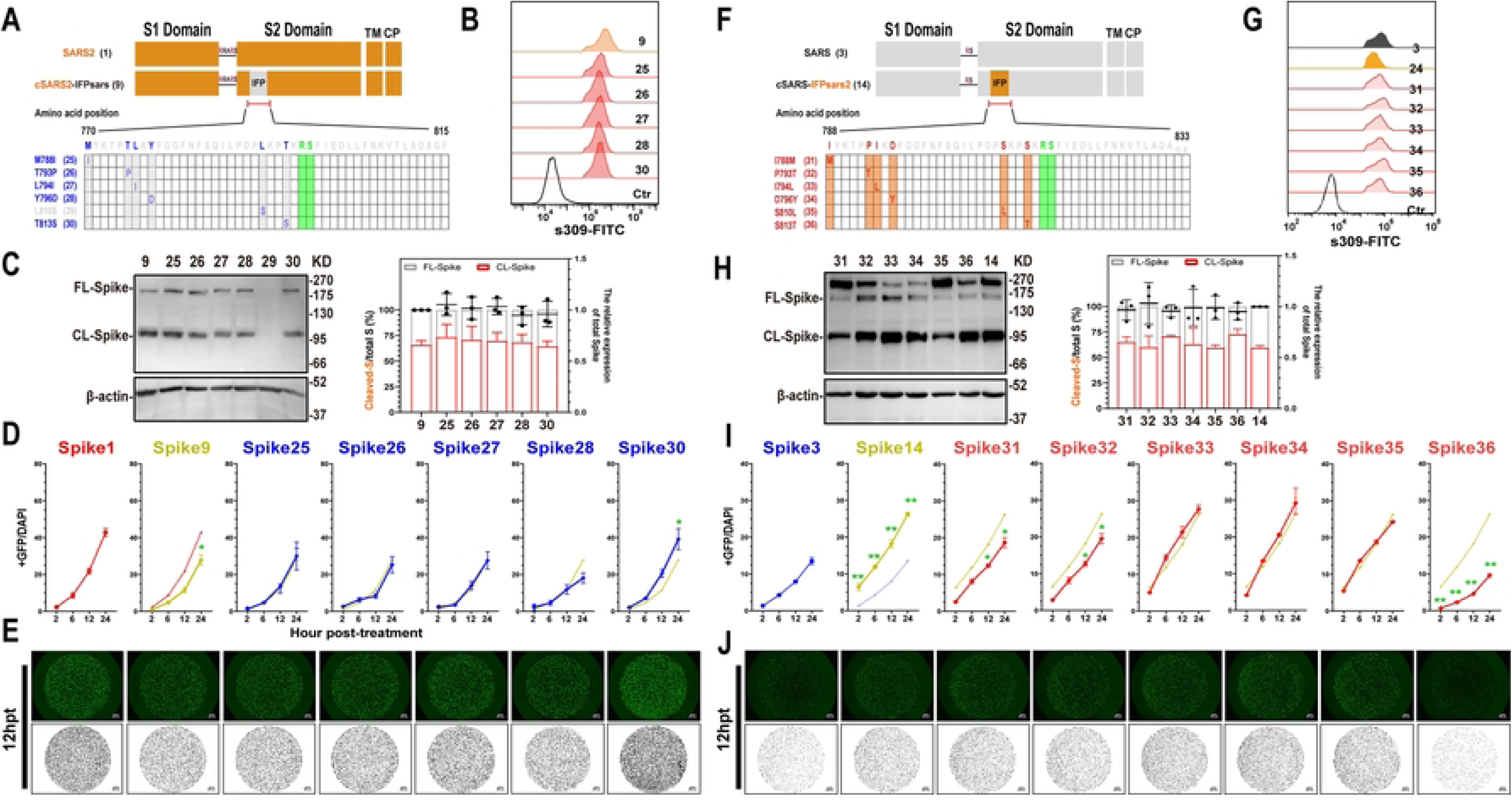
S813T mutation affected the cell membrane fusion ability of IFP-chimeric Spike. (A and F) Schematic diagram of the IFP-chimeric Spike mutants and the numbers in parentheses are identical to those in Figure3B-3D and 3G-3I. Align the sequence of AA 788-833 in SARS2 with SARS (AA 770-815). Residue numbering is shown according to SARS2 S. The mutation sites were marked in red or blue, and the S2’ cleavage sites were in Green. (B and G) FACS. The summarized results of the surface S expression were shown. s309 antibody and mouse anti-human IgG-FITC were used respectively. (C and H) Western blot. Left panel: A representative blot of cell lysates from WT and mutant chimera-Spike expressing 293T cells, FL-Spike and CL-Spike were marked as indicated. Beta-actin was used as a control. Right panel: quantified band intensity using Image J to analyze the protein expression and the ratio of Cleaved-S to the total S. (D and I) Spike-based fusion assay. The fusion activity was quantified by measuring the ratio of GFP+ area to DAPI area by imaging at different times (2, 6, 12 and 24hpt). The results for mutant Spike9 and 14 were shown as yellow-green, Spike25-30 as blue lines, and Spike31-36 as red lines, respectively. (E and J) Representative images of cell-cell fusion. Scale bar: 200 µm. Results are means +/- SD from at least three fields per condition. Results are representative of at least three independent experiments. Statistically significant differences between parental S (Spike9 or Spike14) and mutants were determined by Student’s test at each point (D and I, *: p<0.05: **: p<0.01).

Similar experiments were performed on Spike14. The expressions and cleavages of all mutant, Spike31-36, were comparable to that of the parent S protein, spike14 (Figs 3G and 3H). Despite Spike31 (I788M) and Spike32 (P793T) showing mildly reduced fusion activities, subsequent experiments showed that only the S813T mutation reduced the membrane fusion properties of the S protein significantly (Figs 3I and 3J). These findings corroborated that the mutation of serine to threonine on residue 813 dramatically reduced the fusion ability of chimeric S proteins, suggesting that residue 813 plays a pivotal role in SARS and SARS2 S2 mediated-fusogenicity.

### S813T mutation disturbed the *Sarbecovirus*’s membrane fusion and infection significantly

To further consolidate the above findings, we introduced S813T mutation in S proteins of SARS2 and its variants of concern (VOC) strains (Spike37-41), and T813S mutation in that of SARS. As shown in Figs 4A and 4B, the membrane fusion activities of spikes with S813 were significantly higher than those spikes with T813 in both ACE2-293T and Caco2 cells, and this trend was independent of the native membrane fusion capability of the S protein. The FACS assay confirmed no significant difference in the expression of various S proteins. Even though the levels of S protein cleavage varied between strains, S813 and T813 S proteins of the same strain had roughly the same level of cleavage, with only S813 S proteins of SARS2 and Delta, T813 S protein of SARS having noticeably stronger levels of cleavage than their counterparts. As a result of this finding, it appeared that the promotion of S813 on membrane fusion originated from a mechanism independent of S1/S2 cleavage.

**Fig 4.**
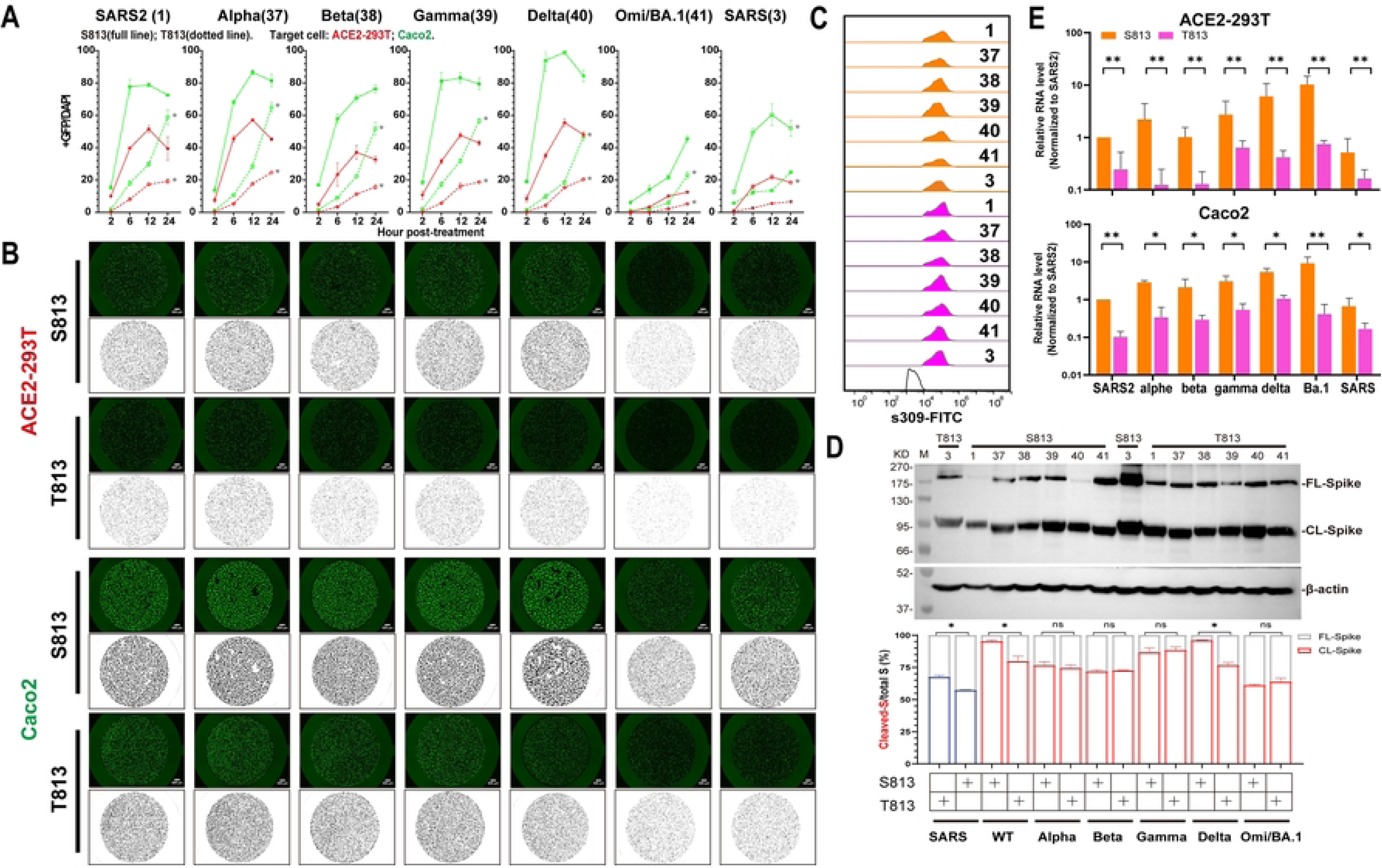
Spike S813T mutation disturbed the membrane fusion and infection of *Sarbecovirus* significantly. (A) Spike-based fusion assay. The fusion activity was quantified at different times (2, 6, 12 and 24hpt). ACE2-293T (Red) and Caco2 (Green) were used, the results of S813 Spike were shown as full lines and T813 Spike as dotted lines. (B) Representative images of cell-cell fusion. Scale bar: 200 µm. (C) FACS. The summarized results of the surface S expression were shown. s309 antibody and mouse anti-human IgG-FITC were used. S813 S and T813 S were shown as saffron and magentas, respectively. (D) Western blot. Top panel: A representative blot of cell lysates from S813 and T813 Spike expressing 293T cells, FL-Spike and CL-Spike were marked as indicated. Beta-actin was used as a control. Bottom panel: quantified band intensity using Image J to analyze the protein expression and the ratio of Cleaved-S to the total S. (E) Pseudovirus assay. The replication of Pseudovirus with S813 or T813 Spike in ACE2-293T and Caco2 cells was determined by RT-qPCR, and the infectivity percentage normalized with that of the virus pseudotyped with Spike1 was shown. S813 S and T813 S were shown as saffron and magentas, respectively. Results are means +/- SD from at least three fields per condition. Results are representative of at least three independent experiments. Statistically significant differences (*: p<0.05) between S813 S and T813 S were determined by Student’s test at each point (A), or two-sided paired t test (D and E).

Considering the pivotal role of S813 in the membrane fusion properties of the S protein, we next interrogated S813 for its possible role in viral infection. We rescued single-cycle infectious SARS viruses by coexpressing the SARS2 replicon [68] and various S genes in package cells. The viruses with various S proteins were used to infect ACE2-293T and Caco2, and the replication products, subgenomic RNAs, were quantified with RT-qPCR. As shown in Fig 4E, the transduction activity of the mutant with S813 S was significantly higher than that of T813 S in SARS2, SARS, or VOC strains. These results indicated that S813T can affect virus entry by modulating the membrane fusion properties of S protein.

### The S813T mutation has no effect on S protein interactions with ACE2

To investigate how S813T mutation alters S fusogenicity, we used a competitive ELISA assay to examine the role of residue 813 in interactions between S protein and ACE2. S813 S and T813 S was separately expressed firstly (Figs 6A). In our study, SARS2 S contained six proline substitutions to generate stabilized and soluble prefusion form [40–42]; SARS S also contained two stabilizing proline mutations in S2 subunit according to an effective stabilization strategy [43,44]. The results showed that S813 S and T813 S proteins had similar affinities towards the ACE2 receptor, indicating that S813T mutation had no effect on S protein receptor binding (Fig 6B). We further verified this finding by assessing the neutralization of RBM-targeting antibodies, which directly blocked the interaction between S protein and ACE2. In this study, we used antibody m396 [45] for SARS, and X65 [15] for SARS2; we found that the representative antibodies showed similar neutralization efficiencies against VSV particles pseudotyped (VSVpp) with parental and mutant S proteins (Fig 5C). In summary, we confirmed that the S813T mutation did not affect S protein interactions with ACE2 and RBD antibodies.

**Fig 5.**
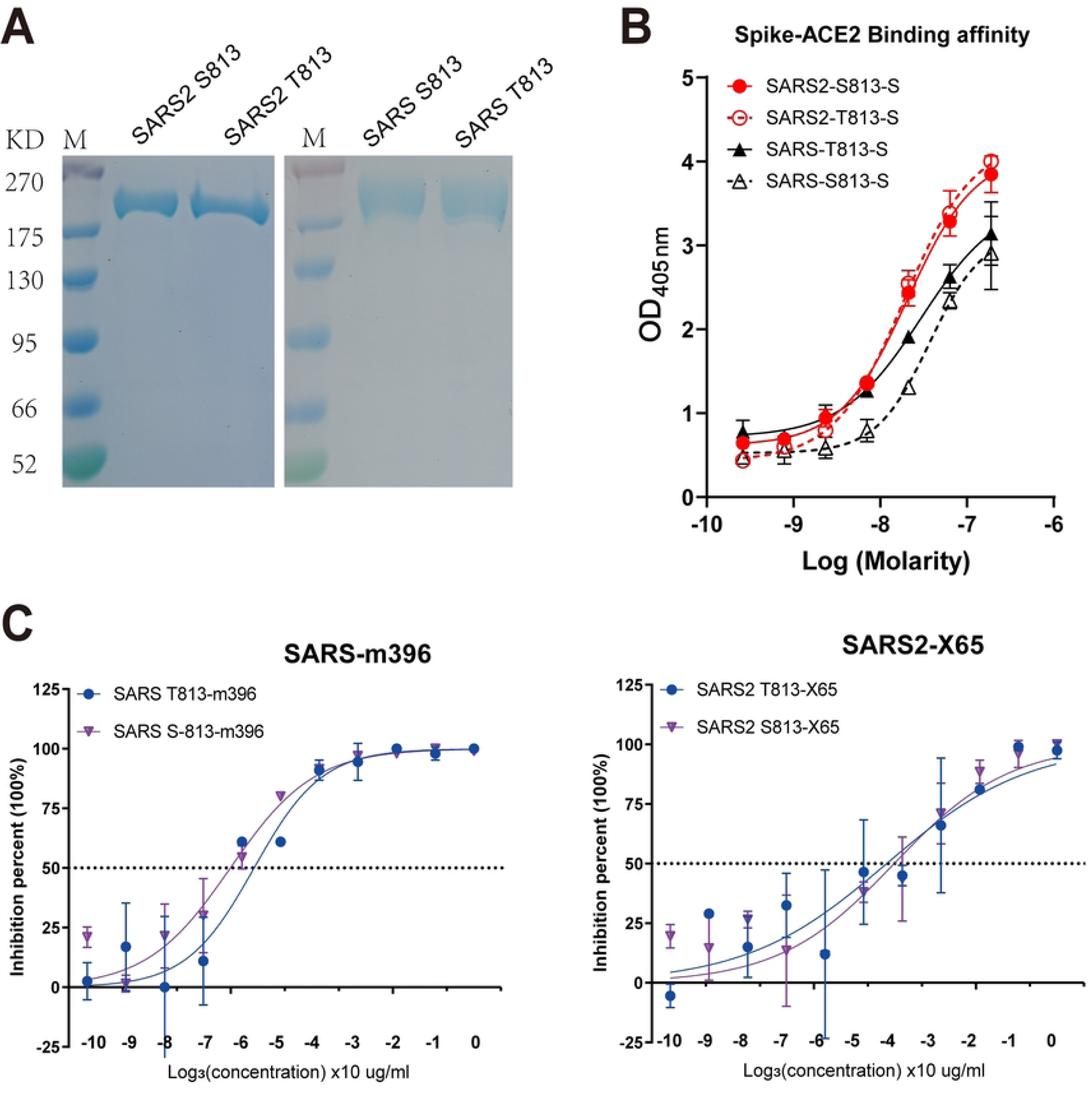
The S813T mutation has no effect on S protein interactions with ACE2. (A) SDS-polyacrylamide gel electrophoresis (PAGE) of SARS2-6P, SARS-2P and Residue 813 substitution S variants. Molecular weight standards are indicated at the left in KD. (B) Competitive ELSIA to detect the binding affinity between S proteins and ACE2. Red line indicated SARS2, and blue line indicated SARS. (C) Neutralization curves for RBD representative antibodies, m396 and X65, with the VSVpp containing parent and mutant S proteins. Each point represents the mean and standard error of 2 independent measurements.

**Fig 6.**
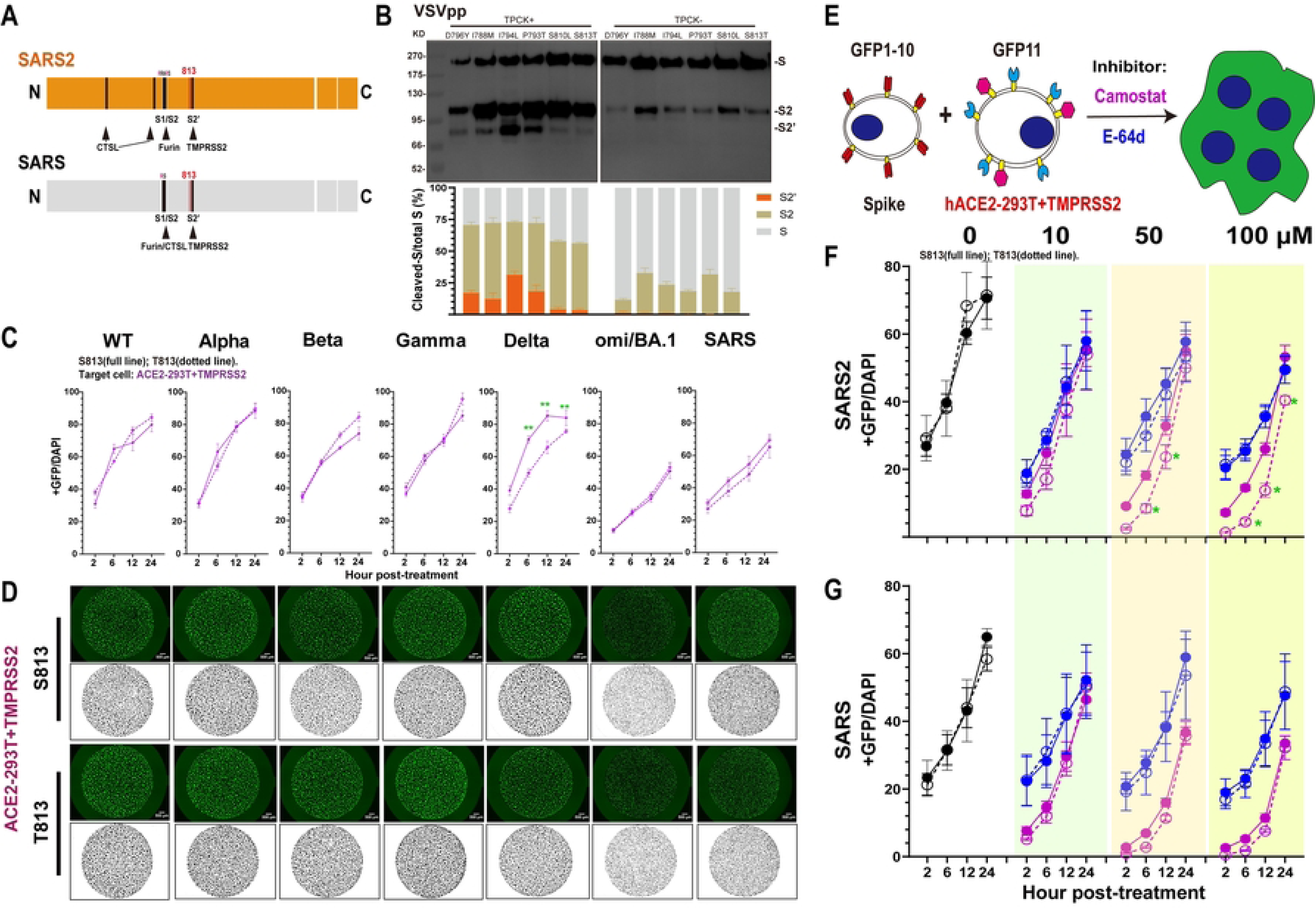
S813T mutation reduced the use of TMPRSS2 by S protein. (A) Schematic illustration of SARS S and SARS2 S including proteolytic cleavage sites: S1/S1, S2’ and CTSL cleavage sites in S1. Residue 813 was indicated as red. Arrow heads indicated the cleavage site. (B) Western blot. Top panel: A representative blot of VSVpp digested with TPCK-trypsin (2 µg/ml at 37℃ for 30 min), FL-Spike (S) and CL-Spike (S2 and S2’) were marked as indicated. Bottom panel: quantified band intensity using Image J to analyze the protein expression and the ratio of S2 and S2’ to the total S. (C) Spike-based cell-cell fusion assay. With the overexpression of TMPRSS2 (purple) in ACE2-293T, the fusion activity was quantified at different times (2, 6, 12 and 24hpt). The results of S813 S were shown as full lines and T813 S as dotted lines. (D) Representative images of cell-cell fusion. The area of cell fusion was shown as green (up) and black (bottom). Scale bar: 500 µm. (E-G) Fusion inhibition of Camostat and E-64d in ACE2-293T+TMPRSS2. A schematic diagram showing the GFP-split system with the inhibitor Camostat and E-64d for Spike-ACE2/TMPRSS2 mediated cell fusion (E). After being pre-incubated with the indicated concentration (0, 10, 50, 100 µm) of Camostat (F) and E-64d (G), the fusion activity was quantified at different times (2, 6, 12 and 24hpt), and the results of Camostat were shown as purple and E-64d as blue. Results are means +/- SD from at least three fields per condition. Results are representative of at least three independent experiments. Statistically significant differences (*: p<0.05) between S813 Spike and T813 Spike were determined by Student’s test at each point (C, F and G).

### S813T mutation reduced the use of TMPRSS2 by S protein

Proteolytic activation of S is critical in the process of CoV entry into cells [46]. It has been reported that efficient infection of SARS and SARS2 requires sequential cleavage of S by furin at the S1/S2 site and then by TMPRSS2 at the S2’ site or by CTSL at two specific sites [47] in endosome when TMPRSS2 expression is repressed (Fig 6A). We noticed that S2’ site is highly conserved in *Sarbecovirus* spikes and residue 813 is located in close proximity to the S2’ site, highly conserved in *Sarbecovirus* spikes, and therefore the 813 change could potentially affect cleavage by TMPRSS2.

To test this hypothesis, we investigated the impact of S813T mutation on the S2’ site cleavage of spike on VSVpp. WB analyses showed that S813T mutation had the most significant effect on S2’ cleavage compared with other point mutations in IFP (Fig 6B). At the same time, we overexpressed TMPRSS2 in ACE2-293T cells firstly (S2 Fig C) and then used the GFP-split system to compare the fusion activities of S813 S and T813 S. We found membrane fusion activities of S813 S and T813 S increased with increasing level of TMPRSS2 expression and both plateaued to similar levels (S2 Figs A and B), this phenomenon was observed for most VOC strains, with only Delta S813 S being stronger than T813 S (Figs 6C and 6D), indicating that increasing TMPRSS2 expression restored the S813T mutation to interfere with S proteins, in other words, the S813T mutation likely reduced S protein sensitivity to TMPRSS2.

To further investigate this possibility, we used the TMPRSS2-specific inhibitor Camostat and examined its effect on membrane fusion under various concentrations, the results (Figs 6E-6G) show that when Camostat concentration was low (10 µM), there was no discernible difference in the ability of S813 S and T813 S in the ACE2-293T/TMPRSS2 system; however, when the Camostat concentration was increased, the difference gradually became apparent, eventually showing that S813 S was significantly higher than T813 S (Figs 6E and 6F). The effect of CTSL inhibitor E-64d was also tested and the results revealed that it had a modest, concentration-independent inhibitory effect in the ACE2-293T/TMPRSS2 system, but did not affect the fusion ability between S813 S and T813 S, suggesting the effect of CTSL on S protein activation was diminished in the presence of TMPRSS2.

Based on these results, it is suggested that the S813T mutation has an effect on S protein fusion pathway modulated by TMPRSS2 by reducing TMPRSS2 cleavage at the S2’ site. S813 S protein had higher utilization and could complete cleavage under lower TMPRSS2 conditions; whereas the T813 S protein used lower and required increased amounts of TMPRSS2 to complete cleavage.

### Evolution AA 813 on Spike in *Sarbecovirus*

Considering the significance of residue 813 in the S protein activation process, we reconstructed an evolutionary tree using the S protein sequences of representative SARS, SARS2 and SARS-relative (SARSr) strains to observe their evolutionary trends in *Sarbecovirus*. At position 813, we observed mostly threonine and serine; threonine was almost only present in two major lineages of SARSr strains, including human SARS-CoV and related viruses; while serine was found in human SARS-CoV-2 strains, bat and pangoline associated SARS-like strains related to SARS2 (such as RaTG13 discovered in 2013 and SL-CoVZXC21 discovered in 2015), as well as a lineage basal to all members of *Sarbecovirus* (Fig 7A). Meanwhile, the mutation frequency of the amino acid at position 813 was calculated with 114 unique SARSr sequences and 10060583 unique SARS2 sequences (GISAID 2020-2022.5.31), respectively. We discovered that in SARS, T813 accounted for 96.49% of the total and S813 accounted for 2.63%; however, in SARS2, S813 accounted for as higher as 99.9%, while T813 accounted for only 0.01% (Fig 7B). Therefore, for SARS-CoV-2 T813 S was likely to be a random mutation that occurred in only a few individuals, without major sustained circulation in human population. Intriguingly, we discovered that other than SARS-CoV and related viruses, the rest of the β-cov, which included highly pathogenic MERS-CoV and SARS2, and the less pathogenic OC43-CoV and HKU1-CoV, were all predominantly serine (Fig 7C).

**Fig 7.**
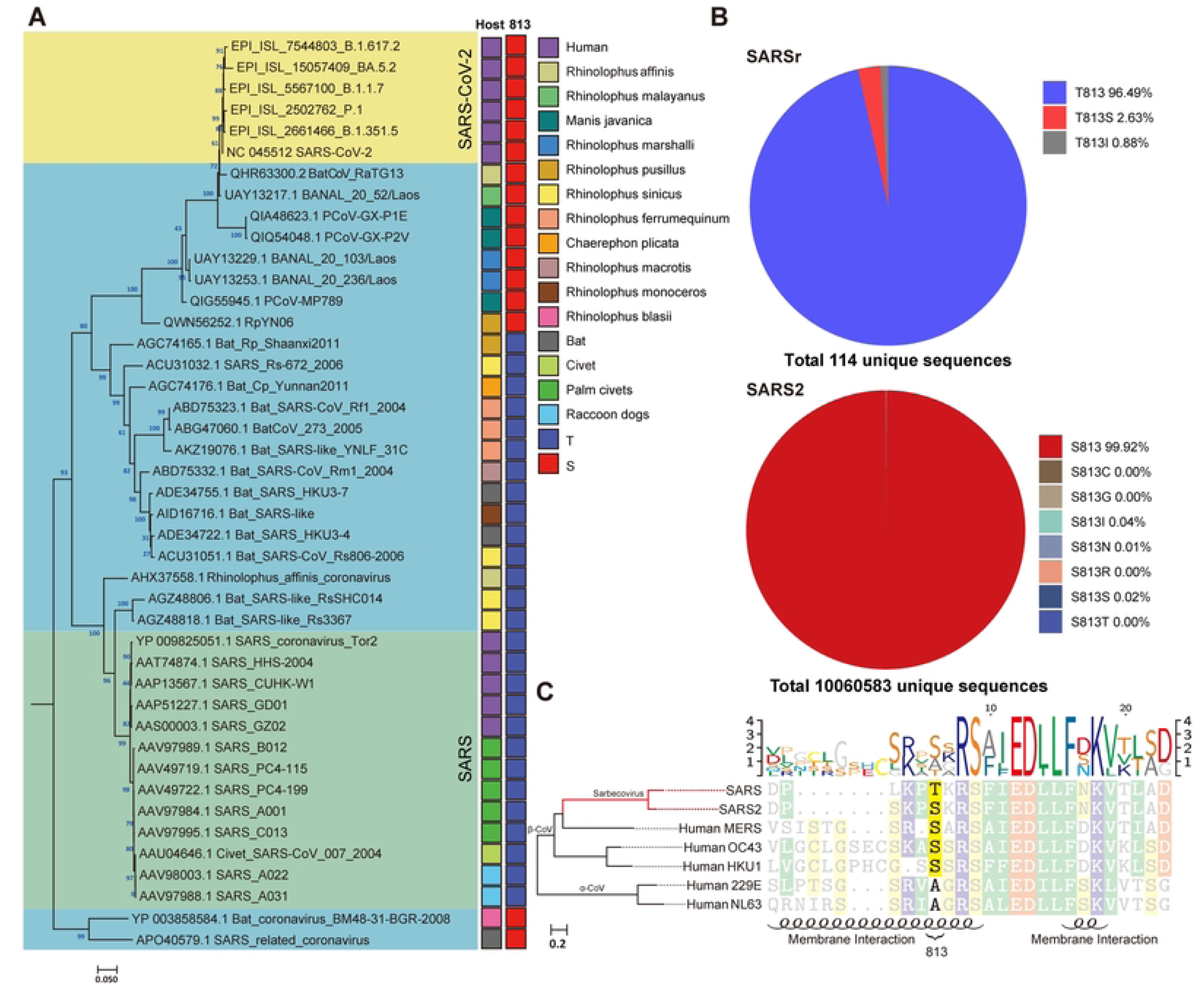

## Discussion

Previous research has proposed that ACE2 binding efficiency [34] and polybasic cleavage site “RRAR” at S1/S2 boundary [31,48] were the primary causes. SARS and SARS2 recognized the same receptor-ACE2 in Humans. The previous study has shown that mutations in the SARS2 RBD make it more accessible to the N-terminal end of ACE2 and stabilize two virus-binding hotspots at SARS2 RBD/ACE2 interface [49]. The polybasic cleavage site of SARS2 has been demonstrated to act as a determinant of transmission [50]. Studies found that the site sensitized SARS2 S protein to fusion-activating proteolysis during virus-cell entry [51]. In this study, we examined the membrane fusion activity of chimeric proteins with various S2s. Our results showed that the activity of S proteins bearing SARS2 S2 is significantly higher than that bearing SARS S2, suggesting that alternation of S2 is sufficient to change the membrane fusion activity of S proteins without disturbing the RBD/ACE2 interaction. We narrowed down the functional domains of S2 to IFP, which is essential for the membrane fusion activity of S protein. Furthermore, we identified the key residue serine 813, as indicated by the data that a single mutation of T813S in SARS S protein could enhance membrane fusion in cell-based membrane fusion assays and viral entry in a single-cycle infectious SARS virus system.

Residue 813 is located immediately upstream of S2’ site (Figure 3A & 3F) and is conserved in various coronavirus strains (Figure. 7C). A recent study proposed that the replication-competent VSVΔG-SARS2 S would especially harbor the S813Y mutation, which reduced the enzymatic activity of TMPRSS2 and increased the stability of S protein for better vaccine design [52]. This unexpected discovery was in agreement with our finding and highlighted the importance of S813 in S protein fusion activity. However, it should be noted that S813Y mutation is not observed in known viral genomic sequences. In contrast, S813 and T813 used in our study are from the SARS2 and SARS sequences, respectively. Thus, our findings were relevant to understanding the real function role of residue 813 during viral entry into the cell.

Host proteases, including furin, TMPRSS2, and CTSL work together to modulate coronavirus S protein-mediated cell fusion [53]. TMPRSS2 is a type Ⅱ transmembrane protein with serine protease activity and is required to trigger cell-cell fusion, it has been reported that TMPRSS2 knock-out 293T cells are unable to form syncytia [54]. TMPRSS2 has been proved to have stronger proteolytic activity against SARS2 S than SARS S. A study has shown that this is due to the multibasic site at S1/S2 boundary, when introduced it into SARS S or ablated it from SARS2 S, the difference can be diminished [36,55]. Here we found that the S2 also affected the utilization of TMPRSS2, the cleavage activity of TMPRSS2 on S813 S was significantly higher than that of T813 S in both SARS and SARS2, and the effect of S813T mutation decreased with increased TMPRSS2 expression. Our data suggested that residue 813, in addition to the multibasic site at S1/S2, could somewhat affect the activity of TMPRSS2 on S protein.

Our study demonstrated that S813T mutation affect the usage of TMPRSS2, did residue 813 impact the function of Furin and CTSL? Furin is ubiquitously expressed in cells [56] and for SARS-CoV-2 S the cleavage site is located in the S1/S2 boundary [54]. Recently, a study found that K814A mutation significantly reduced furin-mediated cleavage of SARS2 S and antagonized pseudovirus transduction, while the S813A had no effect [57]. This work suggests that residue 813 is unlikely to affect furin cleavage. CTSL is a member of the lysosomal cysteine protease and was highly expressed in most human tissues, including the respiratory system, gastrointestinal tract, kidney, and urogenital system [58]. Previous studies have shown that CTSL played a role in the proteolysis of SARS S [59,60] and SARS2 S [26], and the cleavage site of CTSL in SARS S was at (near) S1/S2 [59], while in SARS2 S, it was at two distinct conserved locations in the S1 subunit [47], suggesting residue 813 did not affect CTSL cleavage.

SARS-CoV-2 is a fast-evolving virus, with rapid nucleotide substitution and recombination to generate new strains of altered virulence [61]. A most well-known example was the SARS2 D614G strain rapidly replaced the original virus strain and became the dominant variant [62]. Further studies showed that D614G substitution favored an open conformational state of S protein [63] and promoted syncytium formation through enhanced furin-mediated S cleavage [64]. Through phylogenetic analysis of representative Sarbecovirus S sequences from SARS outbreak in 2002 to the emergence of COVID19 in 2022, we determined that threonine and serine are the only two potential amino acids at position 813 in Sarbecovirus, however, T813S mutation enhanced the S protein membrane fusion function, which might be likely leading to the emergence of SARS-CoV-2 in nature.

In summary, our study demonstrated that residue 813 was a key determinant of S protein fusogenicity and infectivity. The selection and increasing frequency of S813 S following the evolution of Sarbecovirus suggested that the T813S mutation was associated with an improvement of viral fitness through an increased S protein processing and fusogenic potential. These findings have important implications for understanding of the viral S fusogenicity.

## Materials and methods

### Cells and agents

Human embryonic kidney 293T (HEK293T) cells were maintained in Dulbecco’s modified Eagle’s medium (DMEM, Gibco) supplemented with 10% fetal bovine serum (FBS), 100 units/ml penicillin, 100 μg/ml streptomycin, and 2 mM L-glutamine. HEK293T cells that stably express human ACE2 (ACE2-293T) were cultivated in the presence of 2 μg/ml puromycin (Invivogen). Caco2 (human epithelial colorectal adenocarcinoma) cells were cultivated in DMEM (Gibco) containing 20% FBS (TransGen Biotech), 100 µg/ml streptomycin (biosharp), 100 U/ml penicillin (biosharp), 1 mM Non-essential amino acids (NEAA, Gibco). All Cell cultures were incubated at 37°C and 5% CO_2_.

### Plasmids

The pcDNA3.1(+) plasmid backbone was appended with a FLAG-tag sequence (DYKDDDDK) at the C-terminal. The spike coding sequences were all codon optimized for human cells. For the construction of recombinant S plasmids. S2 subunit of SARS-CoV and SARS-CoV-2 VOC strains were amplified by PCR and appended with other regions of the target spike backbone to facilitate In-Fusion cloning.

pQCXIP-BSR-GFP11 and pQCXIP-GFP1-10 were from Addgene (68715,68716); Human TMPRSS2 was amplified from Caco2 cells and cloned into pcDNA3.1(+) with a C-terminal FLAG-tag. All DNA constructs were verified by Sanger sequencing (ACGT).

### Fluorescence-activated cell sorting (FACS)

We conducted FACS using a Beckman CytExpert (Beckman) and data was analyzed with FlowJo software. 293T cells transfected with S proteins for 36h were performed in PBS with 1% BSA. Cells were incubated with primary antibodies on ice for 1h and washed twice with PBS, then incubated with FITC goat anti-human IgG(H+L) (Southern Biotechnology Associates,1:400) for 45 min on ice. Transfection efficiency was assessed by staining with SARS-CoV-2 RBD specific antibodies: s309 [65] (sotrovimab), which had been proved to pan-bind with SARS-CoV-2 VOC strains.

### Western blot analysis

Western blot analysis was performed as previously described procedures [66]. Briefly, cells were washed with ice-cold PBS and soluble proteins were extracted with cell lysis buffer (100mM Tris-HCl pH=8.0, 150mM NaCl, 1% NP-40, phosphatase and protease inhibitor cocktail tablets (Abcam)) according to the manufacturer’s protocol. For the analysis of S protein processing in VSV pseudotyped particles (VSVpp), we loaded 10 ml VSVpp onto 500 μl of a 20% (w/v) sucrose cushion and performed high-speed centrifugation (25000 g for 120 min at 4℃), the concentrated particles were re-suspended in 50 μl PBS. Equal amounts of protein samples were separated by 8% sodium dodecyl sulfate-polyacrylamide gel electrophoresis (SDS-PAGE) and transferred to nitrocellulose filter (NC) membranes. Mouse monoclonal antibodies targeting FLAG tag, β-actin (TransGen Biotech, 1:5000) were used as primary antibodies, horseradish peroxidase-conjugated (HPR) goat anti-mouse IgG antibody (Southern Biotechnology Associate, 1:10000) was the second antibody. The quantitative result of the ratio of cleaved to a full-length spike in immunoblots was analyzed by Image J software.

### Fusion assay

This assay utilized a dual split protein (DSP) encoding the GFP gene, and the respective split proteins, DSP1-10 and DSP11, were expressed in effector and target cells by transfection [38]. For cell-cell fusion activity, hACE2-293T or Caco2 cells transfected with pQCXIP-BSR-GFP11 were prepared as target cells; HEK293T expressing the wild-type (WT) or chimera S proteins and pQCXIP-GFP1-10 were prepared as effector cells. In brief, the 293T cells were grown to 80% confluence in a 12-well plate and transfected with 1 μg pQCXIP-GFP1-10 and 1μg pcDNA3.1(+)-SARS2 S-FLAG (WT or chimera), the hACE2-293T or Caco2 cells in a 12-well plate were transfected with 1μg pQCXIP-BSR-GFP11. After 24h, the target cell and effector cell populations were washed and resuspended in DMEM 10% FBS, mixed at a 4:1 ratio in different combinations, plated at 3x 10^5^ cells per well in a 96-well plate, and the fluorescence images were taken at the indicated time point using a Zeiss LSM800 confocal laser scanning microscope and a Keyence all-in-one Fluorescence microscope BZ-X800. The GFP area was quantified on Image J, and the expression levels of surface S proteins were analyzed using FACS, and the GFP area was normalized to the mean fluorescence intensity (MFI) of surface S proteins, and the normalized values were shown as fusion activity.

### Pseudotyped particles assay

HEK293T cells were transfected with 72 μg WT or chimera S plasmid into a 15 cm cell culture dish. After 24 h, the cells were washed twice and inoculated with VSV*△G-Luc at a multiplicity of infection of 0.1 for 1h. After the inoculum was removed, the cells were washed 5 times using PBS with 2% FPS and further cultured with a DMEM culture medium for 36h. The supernatant was harvested and centrifuged at 3000 rpm to free cellular debris, filtered through a 0.45 μm syringe filter, and stored at – 80°C in small aliquots.

To detect the neutralizing activity of antibodies, serial dilutions (1:3) of mAbs were mixed with an equal volume of 200 50% tissue culture infectious doses (TCID50) SARS2 and SARS VSVpp into a 96 well-plate and incubated for 1 h at 37 °C, and then ACE-293T cells (100 μl, 2 × 10^5^ in DMEM) were added to all wells and incubated for further 24 h at 37 °C. Luciferase activity was analyzed by the luciferase assay system (Promega). IC50 was determined by a four-parameter logistic regression using GraphPad Prism 8.0 (GraphPad Software Inc.)

### Proteins and Monoclonal antibodies expression and purification

Proteins and antibodies were generated as described previously [67]. In brief, for S proteins and ACE2, target genes were firstly amplified and subcloned to pcDNA3.1(+) vetor, and after performing Site-Directed Mutagenesis, plasmids were transfected into HEK293T cells using polyethylenimine (PEI) and cultured for 5 days. The supernatant was collected and purified using Ni-sepharose. For antibodies, the single B cells of COVID19 convalescents were obtained and then sorted into 96-well plates, and the IgG heavy and light chain variable genes were amplified by reverse transcriptase polymerase chain reaction (RT-PCR) and cloned into human IgG1 expression vectors and co-transfected into HEK293T cells with equal amounts of heavy/light-chain plasmids. Five days post-transfection, the supernatants were collected and purified using protein A agarose beads.

### Rescue and infection of recombinant SARS-CoV/SARS-CoV-2 virus

2×10^6^ HEK293T cells were seeded in a 6-cm plate. After recovery for 24 h, HEK293T cells were transfected with three µg Rep (ref) and three µg plasmids encoding various S genes using Hieff TransTM Liposomal Transfection Reagent (Yeasen Biotech, Cat#40802ES03, Shanghai, China). Six h post-transfection, the medium containing the mixture of DNA/transfection reagent was replaced with fresh medium. After recovery for 36 to 48 h, the supernatants were collected for further use.

2×10^5^ HEK293T-ACE2 cells were seeded in one well of a 6-well plate. After recovery for 24 h, the HEK293T cells were infected with recombinant SARS-CoV/SARS-CoV-2 viruses, and two h post-infection, the viruses-containing medium was replaced with fresh medium. 24 h post-infection, the cells were collected for the extraction of total RNA and/or protein.

### Enzyme-linked immunosorbent assay (ELISA) to detect Spikes binding ability to ACE2

ELISA was performed as described previously [67]. High-protein binding microtiter plates (Costar) were coated with 2 μg/ml human ACE2 protein in PBS overnight at 4℃ respectively. After 3% BSA in PBS blocking, serially diluted S proteins 1:3 starting at 50 ng/μl were incubated for 1h at 37℃. After washing 6 times with PBST, a S2 antibody from our lab, I24, was incubated at 10 μg/ml at 37℃ for 1h, after washing again, the HPR-conjugated goat anti-human IgG antibody (Jackson Immuno Research, 1:2000) was incubated for another 1h at 37℃. The plate was developed with Super Aquablue ELISA substrate (eBiosciences). Absorbance was measured at 405 nm on a microplate spectrophotometer (BioTek).

### Effect of drug treatment on fusion ability and cell viability assay

Camostat and E64d were diluted at different concentrations first and then added to the cell mixture for fusion assay in a 96-well plate. The cell mixture was then mixed gently and cultured in a 5% CO_2_ environment at 37℃ for subsequent testing.

The effects of Camostat and E64d on cell viability were measured by CCK8 assay. 293T cells were seeded into a 96-well plate and were left untreated or treated with different concentrations of drugs for 24 h. After treatments, CCK8 was added into the culture medium and incubated for 1 h at 37 °C to measure the absorbance at 405 nm.

### Phylogenetic analyses

We selected representative sequences from the NCBI taxonomy *Sarbecovrius* grouping and compared them using mafft software. We then used fasttree software to construct a phylogenetic tree based on SARS-like S proteins.

### Statistical analysis

The Prism software (Graphpad Version 8.0) was used for all statistical analyses. The significance of differences between the two groups was determined with a two-tailed Student’s t-test. One-way or two-way analysis of variances with Bonferroni correction was employed for multi-group comparison. For all analyses, only a probability (*p*) value of 0.05 or lower were considered statistically significant (p > 0.05 [ns, not significant], p % 0.05 [*], p % 0.01 [**], p % 0.001 [***]).

## Supporting information

**S1 Fig. Reversing IFP motif of S2-chimera Spike influenced the fusogenicity significantly.** (A and F) Schematic diagram of the S2 motif chimera Spike. The S2 was divided into 3 parts as Figure2 and to replace the corresponding area in turn. The F1 was further divided into two parts just as Figure2 did. The numbers in parentheses are identical to those in Figure3B-3E, 3F-3J. (B and G) Western blot. A representative blot of S-expressing cells (top) and quantified band intensity (the ratio of CL-S to the FL-S plus CL-S proteins) (bottom) are shown. (C and H) FACS. The summarized results of the surface S expression were shown. s309 antibody and mouse anti-human IgG-FITC were used respectively. (D and I) Spike-based fusion assay. The fusion activity was quantified by measuring the ratio of GFP+ area to DAPI area by imaging at different times (2, 6, 12 and 24hpt). The results for SARS2, Spike4, 15-19 or SARS, Spike2, 10-14 were shown as Saffron and grey lines, respectively. (E and J) Representative images of cell-cell fusion. Scale bar: 500 µm.

Results are means +/- SD from at least three fields per condition. Results are representative of at least three independent experiments. In B and G, statistically significant differences between parental S (Spike2 or Spike4) and chimeric Spikes were determined by a two-sided paired t test (*: p<0.05). In D and I, statistically significant differences between parental S (Spike2 or Spike4) and chimeric Spikes were determined by Student’s test at each point (*: p<0.05: **: p<0.01).

**S2 Fig. The Influence of TMPRSS2 on Spike fusogenicity.** (A) Spike-based fusion assay. The fusion activity was quantified by measuring the ratio of GFP+ area to DAPI area by imaging at different concentration of TMPRSS2 (0, 0.5, 1 and 2 µg). The results for SARS2 and SARS were shown as Red and grey lines, respectively. (B) Representative images of cell-cell fusion. Scale bar: 500 µm. (C) Western blot. A representative blot of 293T cell lysates expressing TMPRSS2 with various concentrations. Beta-actin was used as a control. (D) Cell viability assay. The cell viability with different doses of Camostat and E64d was evaluated by CCK8 assay.

Results are means +/- SD from at least three fields per condition. Results are representative of at least three independent experiments. Statistically significant differences (**: p<0.01) between S813 Spike and T813 Spike were determined by Student’s test at each point (C, F and G).

**S3 Fig. Genomic description of AA 813 in Spike of *Sarbecovirus*.** (A) The phylogenetic tree based on *Sarbecovirus* S proteins (SARS-like strains, n=24 genomes; SARS strains, n=13 genomes; SARS2 strains, n=6 genomes). All strains invariantly containing serine at position 813 were marked red while containing threonine were marked mazarine. Color coding as indicated according to species. (B) Amino acid frequency of site 813. 114 complete spike protein sequences of SARS were collected from NCBI and 10,060,583 complete spike protein sequences of SARS2 (2020-2022) were collected from GISAID. The method of analysis was performed as previously described [69]. Briefly, After removing redundant sequences with 100% sequence identity and multiple sequence alignment (MSA), site 813 based on the reference sequence of SARS2 was derived and the amino acid frequency of site 813 can be calculated based on the un-redundant dataset of SARS and SARS2, respectively. (C) Phylogenetic tree of Human coronavirus. The representative strains of 7 human coronaviruses were clustered by amino acid sequence phylogeny and observed the diversity of AA 813. In α-CoV (229E and NL63), there was only Alanine, while in β-CoV, Serine was in SARS2, MERS, OC34 and HKU1; threonine only in SARS. We used the WAG+F+I+G4 optimal model of the Iqtree software to construct a phylogenetic tree based on S proteins. The right-hand sequence mapping was based on texshade software for mapping. The secondary structure, i.e. the membrane fusion region, was predicted using the PSIPRED web page.

## Acknowledgements

We would like to acknowledge Dr. Wenxia Tian at College of Veterinary Medicine, Shanxi Agricultural University for providing materials, technical support and valuable suggestions. We thank Dr Hai Li at School of Basic Medical Sciences, XI’AN JIAOTONG University for suggestions and revision of the manuscript.

## Author Contributions

**Conceptualization:** Yong Ma, Yao-Qing Chen.

**Data curation:** Yong Ma, Tianyi Qiu, Pengbin Li.

**Formal analysis:** Yong Ma, Pengbin Li, Yunqi Hu.

**Funding acquisition:** Yao-Qing Chen, Yong Ma.

**Investigation:** Yong Ma, Pengbin Li, Yunqi Hu, Tianyi Qiu, Wenjing Zhao, Jun Chen,Xiaoli Xiong, Mang Shi, Ji-An Pan3, Yao-Qing Chen

**Methodology:** Yong Ma, Pengbin Li, Yunqi Hu, Tianyi Qiu, Mang Shi, Jiaxin Zhuang, Ji-An Pan.

**Project administration:** Yao-Qing Chen.

**Resources:** Hongjie Lu, Kexin Lv, Mengxin Xu, Jiaxin Zhuang, Xue Liu, Bing He, Shuning Liu, Lin Liu, Yuanyuan Wang, Xinyu Yue, Yanmei Zhai, Wanyu Luo, Haoting Mai, Jun Chen.

**Software:** Tianyi Qiu, Mang Shi.

**Supervision:** Yao-Qing Chen.

**Validation:** Yong Ma, Pengbin Li, Yunqi Hu.

**Visualization:** Yong Ma, Tianyi Qiu, Xue Liu, Lixiang Wang, Mang Shi.

**Writing – original draft:** Yong Ma, Pengbin Li.

**Writing – review & editing:** Yong Ma, Xiaoli Xiong, Mang Shi, Yao-Qing Chen.

